# An alphabet of allostery: a transferable transfer entropy contact patterns predicts allosteric-site character and conformational rewiring

**DOI:** 10.64898/2026.07.29.741458

**Authors:** Aysima Hacisuleyman

## Abstract

Allostery is increasingly understood as the propagation of dynamic information through a protein, yet the computational descriptors of that flow are computed one structure at a time and carry no transferable, sequence-level prior. Here we build an alphabet of allostery: a dictionary of local contact words, short sequence windows anchored by three-to four-residue spatial cliques, each carrying a distribution of net Gaussian network model transfer entropy scores pooled over a non-redundant set of Protein Data Bank structures. The alphabet consists of 131,611,766 unique words drawn from 212,860,934 clique observations. Projecting any protein’s sequence and structure onto this dictionary yields a per-residue allosteric track, net TE, source, sink, and switch channels, with no system-specific fitting. Validating against the Allosteric Database, annotated allosteric-site residues behave as transfer entropy sinks (information receivers): the sink channel discriminates sites from the rest of the protein with pooled ROC-AUC = 0.543 over 646,629 residues (permutation z = 19.1), an effect small in magnitude but overwhelmingly significant and robust to word-frequency leakage. The directional channels are mechanistically informative in a two-state experiment on nine canonical allosteric proteins: source residues predict the largest apo→holo conformational rewiring (meta-analytic Spearman ρ = +0.106, positive in 7/9 proteins) while sink residues mark the most conformationally stable positions (ρ = −0.105, 8/9). Sinks thus mark where allosteric signal is received and sources mark where it drives motion. Finally, we mine the most context-variable words into a compact, hydrophobic-enriched switch vocabulary that we propose as a design dictionary for engineering allosteric mechanisms.

## 1. Introduction

Allostery is the functional coupling of distant sites in a protein.^1, 2^ It is a mechanism by which enzymes are regulated.^3^ A modern view of allostery treats it as a dynamic information flow^1, 2, 4^, where perturbations at one site propagate through a protein’s fluctuation network to modulate distant regions, even without large conformational changes.^4–6^ Since a single static structure cannot capture this mechanism^7, 8^, predicting multi-state ensembles and conformational transitions remains a central challenge for the field.^9^ To address this, elastic network models (ENMs), particularly the Gaussian network model (GNM), offer a computationally efficient framework to capture these fluctuations and collective motions.^10–12^ By computing the transfer entropy(TE) over GNM^13–15^, we can establish a directional measure of which residues drive, and which respond to, collective motion.

Yet the limitation of TE is transferability. It is computed per structure and per conformation, and the resulting per-residue scores live only in the coordinate frame of that structure. Sequence-based allostery predictors exist^16–19^, but they are trained end-to-end on-site annotations rather than built from the physics of information flow.

Here we ask a different question: is there a transferable pattern of local contact motifs that carry allosteric signal across proteins? To answer it constructively, we compute the GNM derived transfer entropy across a culled, non-redundant PDB dataset generated using PISCES^20^, decompose each structure into local contact words anchored by spatial cliques, and pool the per-word TE scores into an alphabet. Each word is recorded with their distribution, mean, spread, and frequency, of the net TE experienced by its anchoring residues across every protein in which the motif appears. The alphabet then becomes a reusable lookup table: any new sequence with a structure can be projected onto it to yield a per-residue allosteric track, with no fitting.

This work makes four contributions. Building the alphabet itself, a pooled, frequency-weighted dictionary of 131.6 M *contact words*, and releasing a scoring method that maps it onto any PDB chain to produce the net TE, source, sink, and switch tracks. Using this method, we established a validated directional signal at allosteric sites. Using the Allosteric Database (ASD)^21^ as the ground truth, annotated allosteric-site residues are found to be transfer entropy *sinks*, with an effect that is small but significant and robust to word-frequency leakage. We then provide a mechanistic decomposition of the directional channels through a two-state apo → holo experiment across nine canonical allosteric proteins, in which source and sink residues play opposite and complementary roles. Finally, we mine the most context-variable words into a compact, hydrophobic-enriched switch patterns of graftable coupling motifs, offered as a design dictionary.

## 2. Methods

### 2.1 GNM and transfer entropy

#### 2.1.1 Structures and elastic-network representation

Each protein chain is represented as a graph whose nodes are residues and whose edges join residues that are in contact. For each residue the contact coordinate is its centroid, the mean position of all its atoms. A residue pair is connected when the distance between their centroids falls below a cutoff *r_c_*. Since transfer entropy estimates over the Gaussian network become unstable for large, sparsely sampled chains, *r_c_* is size-dependent: 7.5 Å for chains shorter than 250 residues and 9.5 Å for chains of 250 residues or more. The resulting contact Kirchhoff matrix, *Γ*, has *Γ* □ *=* −1 for contacting pairs (*i* ≠ I ≠ *j*) and *Γ* = −*Σ* □ ≠ *Γ* □ on the diagonal.

#### 2.1.2 Transfer entropy

We quantify the directed allosteric communication by the transfer entropy, *T*_*i→j*_, computed from the dynamic Gaussian Network Model (dGNM) of Hacisuleyman and Erman^15^, built on Schreiber’s^22^ transfer entropy concept, which measures how much the past fluctuations of residue *i* reduce the uncertainty about the future fluctuations of residue *j*. The dGNM statistics follow from the eigen decomposition of *Γ*: with non-zero eigenvalues *λ*_*k*_ and eigenvectors *u*_*k*_, the equilibrium cross-correlation is 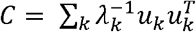. From the directional pair *T*_*i→j*_ and *T*_*j→i*_, we construct the net transfer entropy (net TE) on each pair,

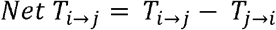

an antisymmetric quantity that is positive when residue *i* is the information source for residue *j*. Summation of the outgoing net TE at each residue gives the signed per-residue score used throughout, *Net TE = Σ □ (T*_*i→j*_ *-T*_*j→i*_), positive for sources and negative for sinks.

The transfer entropy is evaluated at a single characteristic delay *τ*, chosen automatically from each structure’s own GNM dynamics rather than a default value: for every residue we compute the normalized autocorrelation of its position fluctuations, 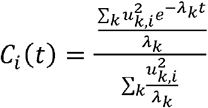 (sum over the non-zero GNM modes k), and define its relaxation time as the time t at which *C*_*i*_ (*t*) decays to 1/e of its initial value. The single *τ* used for the whole chain is the median of these per-residue relaxation times, which makes the delay a robust, parameter-free property of each structure’s own dynamics.

The signed per-residue *Net TE*, together with its source and sink parts, is the quantity projected onto the contact-word alphabet in Section 2.2.

### 2.2 Alphabet construction

From each chain we enumerate spatial cliques (three to four fully connected sets of residues within 8.0 Å apart from each other). Each clique defines a *word*: the sequence window spanning its extreme residues, with clique members upper-cased and linkers lower-cased, derived from our previous study^23^. Each word occurrence carries the *Net TE* of its clique residues. Words are pooled over the PISCES^20^ set, a non-redundant precompiled PDB list; for each distinct word we accumulate the *mean, standard deviation*, and the *count* of its *Net TE* values. The pooled database (131,611,766 words) is stored as Parquet^24^ and queried via DuckDB^25^.

### 2.3 Sequence-scoring method

The sequence scoring method exposes *Reference(parquet, min_count)* (a DuckDB-backed word → (mean, std, count) lookup) and *AlloAlphabet.score_chain(structure, chain)*, which enumerates a chain’s words, looks up their statistics, and assigns each residue a log-count-weighted mean (*weight log*_*10*_*(count) + 1*) over all covering words, producing the *Net TE*/ *source* / *sink* / *switch* / *n_words* channels. *write_scored_pdb* writes any channel into the B-factor column.

### 2.4 ASD validation

ASD(2023 release)^21^ allosteric-site residues were parsed to per-chain residue labels. Each scored residue was z-normalized within its chain. Discrimination was measured as ROC-AUC (analytically, *AUC = U / (n1n0*) from the Mann–Whitney *U*), PR-AUC (average precision from descending-score cumulative precision/recall), per-chain median AUC, and top-quantile enrichment. Significance was assessed by a within-chain label-permutation null (200 permutations preserving per-chain site counts), reporting the *z* of the observed pooled AUC against the null. Leakage robustness was assessed by repeating at a word-frequency floor of count ≥ 30.

### 2.5 Two-state (apo → holo) experiment

Nine allosteric proteins with two states were selected. For each, per-residue *Net TE* was computed in both states, z-normalized within each state, and the rewiring defined as |Δ *Net TE*| over residues common to both structures (matched by residue number). Each alphabet channel (evaluated on the apo state) was correlated with |Δ *Net TE*| by Spearman ρ, pooled and per-protein. Per-protein ρ values were combined by sample-size-weighted Fisher-z meta-analysis, reporting the meta ρ and 95% CI.

### 2.6 Switch-vocabulary mining

Words with *Net TE* standard deviation above mean + ~1.95 SD of the common-word (count ≥ 30) distribution (threshold 1.945) were designated switch candidates. They were characterised by mean-TE sign bias, linker-length distribution, and per-position amino-acid enrichment (log_2_ fold-change vs background), and ranked by *design scores = standard deviation* × *(log*_*10*_ *count* + 1).

## 3. Results

### 3.1 An alphabet of contact words

We define a *contact word* as the short sequence window anchored by a spatial clique, a set of three or four residues all in mutual contact with their centroid distance ≤ 8.0 Å. The word is the linear sequence spanning the extreme clique residues; clique members are written in uppercase and intervening linker residues in lowercase (e.g. ClAS, RfLR). Each word inherits, from the structure it was found, the net GNM TE values of its anchoring residues. Pooling over the PISCES^20^ set, we accumulate for every distinct word the distribution of net TE values it experiences across all its occurrences.

The resulting alphabet contains 131,611,766 unique words from 212,860,934 clique observations, in two letter classes: approximately 78.3 M three-residue cliques and 53.3 M four-residue cliques. Word length spans from 3 to over 1,500 residues (reflecting long-range contacts), but the mass of the distribution is at short windows (median linker-inclusive length ≈ 4). The alphabet is heavy tailed in frequency: 86.7% of words are singletons. Analyses use words with count ≥ 2 by default, and a common-word subset (count ≥ 30; 306,658 words) for statistics that must be robust to single-structure irregularities (Figure 1).

**Figure 1.**
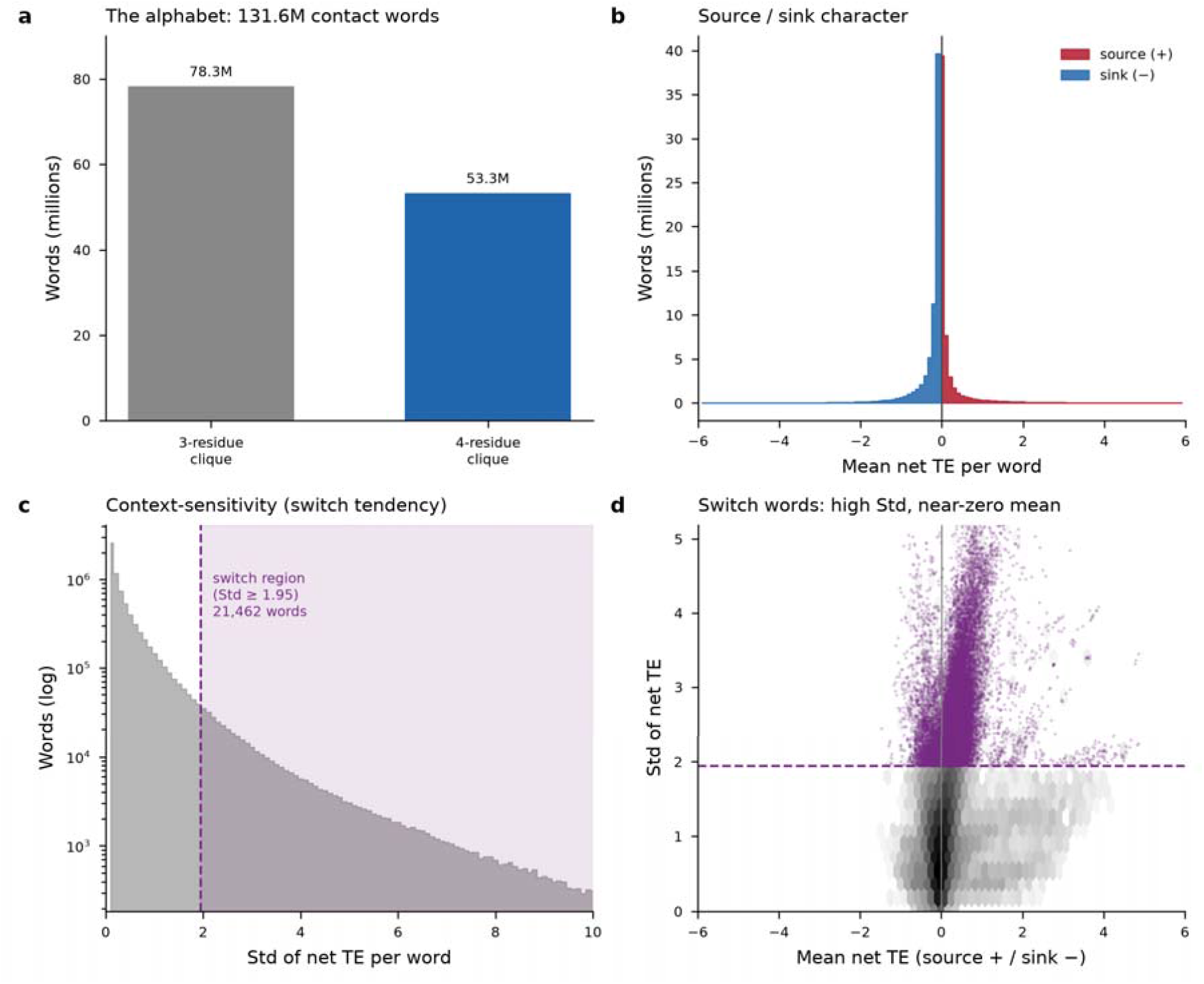
The alphabet of contact words: clique-class composition (a), source/sink character of per-word mean net TE (b), the context-sensitivity (switch) distribution with the switch band marked (c), and switch words in the mean–Std plane (d).

For each word we retain three summary statistics: the *mean net TE* (positive = net source/emitter, negative = net sink/receiver), the *standard deviation* of net TE across occurrences (how context dependent the word’s dynamic role is, the “switch” channel), and the *occurrence count* (used for frequency weighting). The switch threshold, set at the mean + ~1.95 SD of the common-word standard-deviation distribution, is 1.945; 21,462 words exceed it and constitute the switch-candidate set analyzed in Section 3.5.

### 3.2 A per-residue allosteric track for any sequence

The scoring method projects the pooled alphabet onto any structure. For a given PDB chain it enumerates every contact word present, looks up each word’s pooled statistics in a DuckDB^25^-backed index, and assigns to every residue a log-count-weighted mean over all words in which it participates (weight = log_10_(count) + 1, so that frequently observed words dominate). This yields five per-residue channels: net_te (signed log-count-weighted mean net TE), source (max(0, net_te), the emitter component), sink (max(0, −net_te), the receiver component), switch (log-count-weighted mean of the per-word TE standard deviation), n_words (coverage (number of words covering the residue).

As a first illustration, we apply this to a protein whose allosteric architecture is textbook-characterised. On adenylate kinase (PDB id: 1AKE:A)^26^ the method achieves 100% residue coverage, and the mobile LID (residues 118–167) and NMP-binding (30–67) subdomains, the enzyme’s canonical allosteric elements, show elevated source and switch scores (Figure 2), a first sanity check that the alphabet localizes dynamically active regions. The *write_scored_pdb* function writes any channel into the B-factor column for direct rendering in molecular viewers.

**Figure 2.**
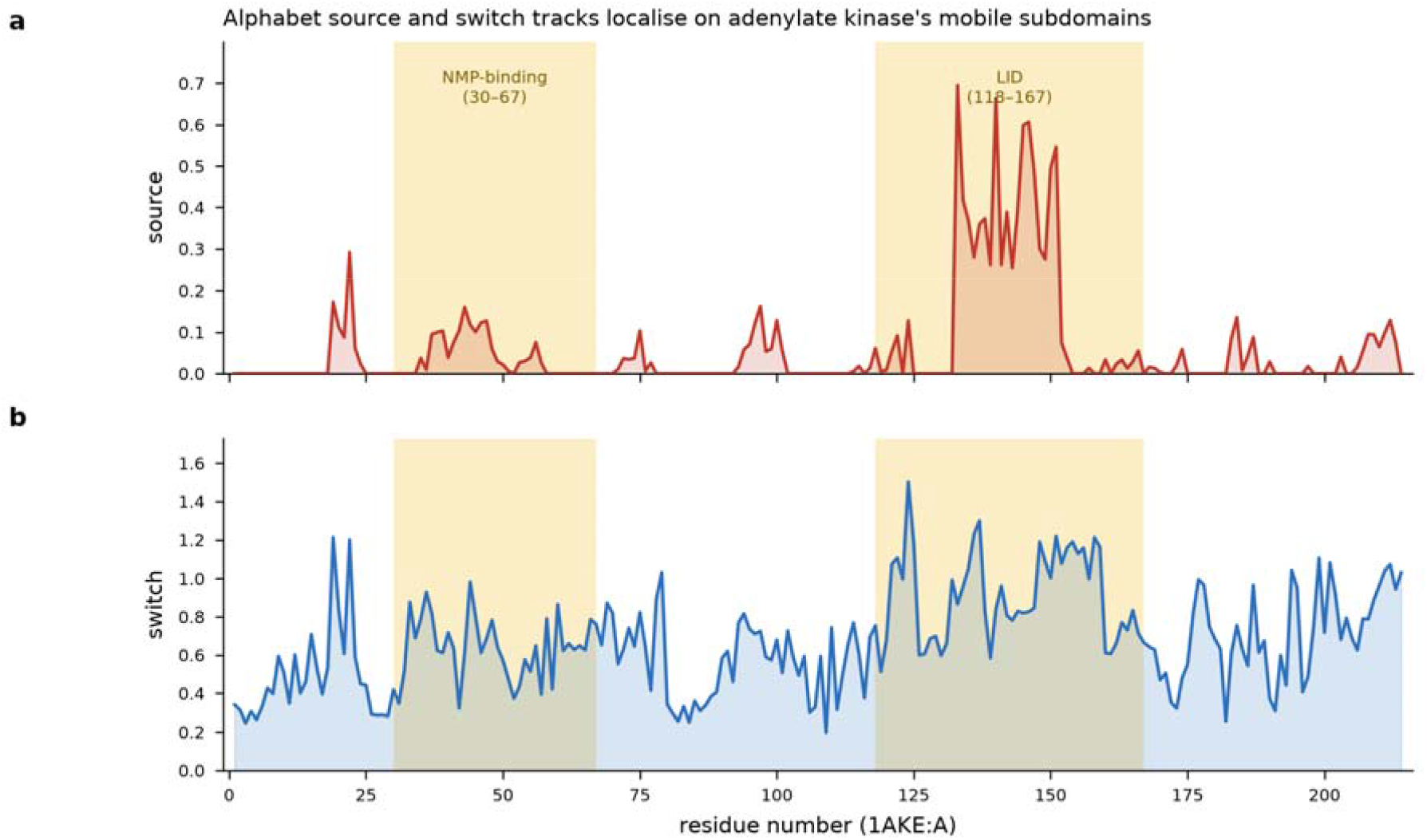
Per-residue alphabet tracks on adenylate kinase (1AKE). The source (a) and switch (b) channels are elevated over the mobile NMP-binding (residues 30–67) and LID (118–167) subdomains (shaded), the enzyme’s canonical allosteric elements, illustrating that the pooled alphabet localizes dynamically active regions on a single structure.

### 3.3 Allosteric sites are transfer entropy sinks

To test whether the alphabet carries real allosteric signal, we validated it against the ASD(2023 release)^21^, which annotates experimentally characterized allosteric sites. We scored 1,773 chains across 1,391 PDB entries (646,629 residues; 19,101 annotated allosteric-site residues; 99.4% residue coverage). Since each protein has its own TE baseline, every residue’s scores were z-normalized within its own chain before site-versus-non-site discrimination was measured, both pooled and per-chain. ROC-AUC and PR-AUC were computed analytically from the Mann– Whitney *U* statistic (no model fitting is involved). The result is directional and clear: the sink channel is the discriminator (Figure 3, Table 1).

**Table 1.** Discrimination of ASD allosteric-site residues by alphabet channel.

| Track | Pooled AUC<br>(count $\geq 2$) | PR-AUC<br>(base 0.030) | Per-chain<br>median AUC | Top 10%<br>enrichment | Permutation z |
| --- | --- | --- | --- | --- | --- |
| sink | 0.543 | 0.034 | 0.525 | 1.23× | 19.1 |
| absolute net-TE | 0.483 | 0.029 | 0.480 | 1.01× | -7.1 |
| switch | 0.494 | 0.029 | 0.500 | 0.86× | — |
| net-TE | 0.452 | 0.026 | 0.468 | 0.85× | — |
| source | 0.436 | 0.026 | 0.455 | 0.77× | — |

**Figure 3.**
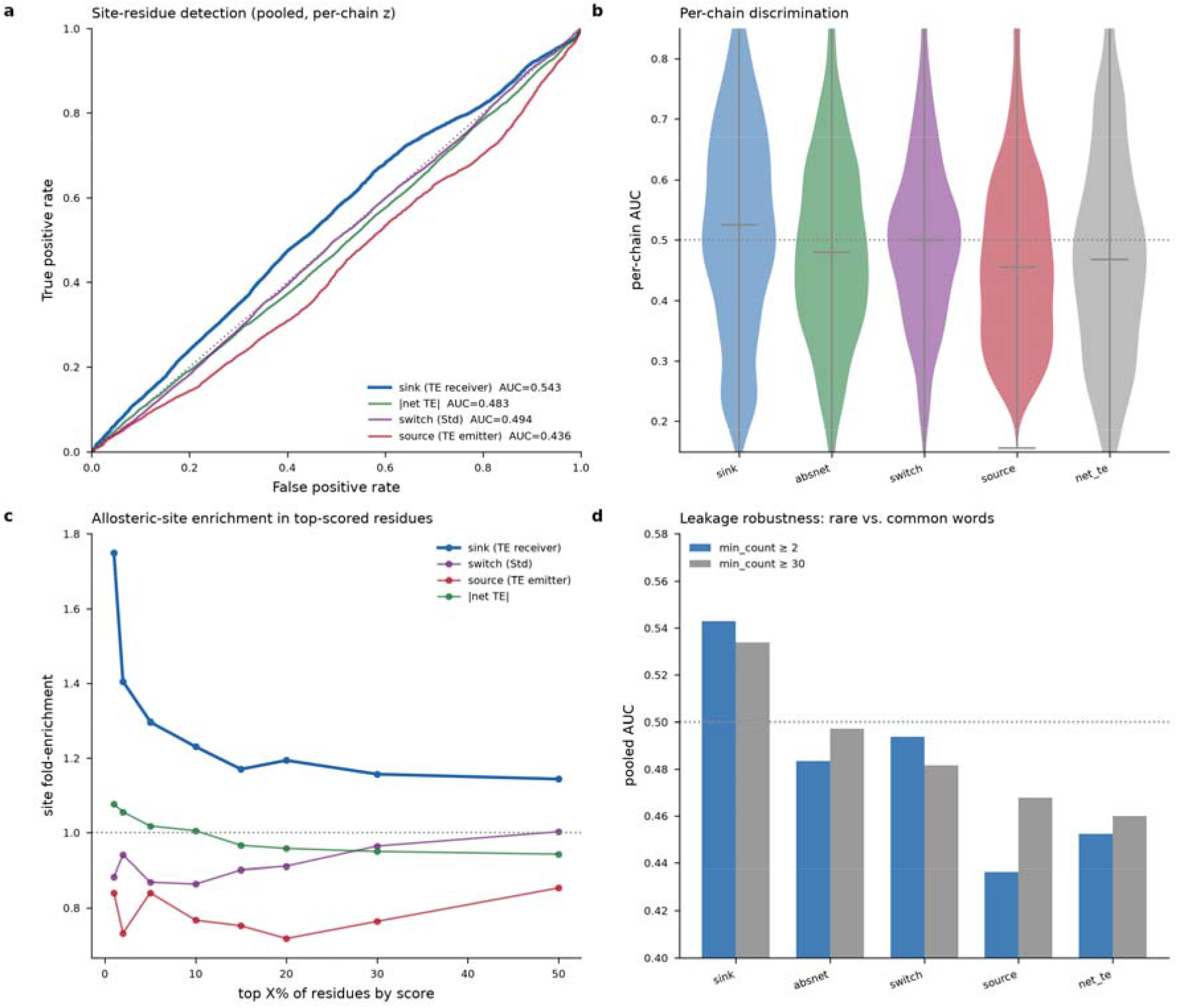
ASD validation. ROC by channel (a), per-chain AUC distribution across 1,763 chains (b), allosteric-site enrichment in top-scored residues (c), and leakage robustness across word-frequency floors (d).

Sink score is positive-discriminating in ~55% of the individual chains, and ASD sites are enriched 1.23× in the top-scored quantile. The source and signed net TE channels are anti-predictive (AUC 0.44–0.45, equally far from chance but with inverted sign, high-source scoring residues are depleted, not enriched), so the effect is genuinely about *receiving*, not merely experiencing, allosteric information.

Two controls establish that the signal is real rather than an artefact of the pooling. First, a within-chain label-permutation null (200 permutations preserving each chain’s site count) places the observed pooled sink-AUC at *z* = 19.1 above the null, overwhelmingly significant given the 646,629-residue sample. Second, since a word seen in few structures could in principle lead to memorization of a training protein, we repeated the analysis restricted to common words (count ≥ 30), which caps any single structure’s contribution to ≤ 3.3%. The median single-structure leakage fraction was 14%, so this control was necessary, and the sink signal survives essentially intact (AUC 0.534, z = 15.3).

An AUC of ~0.54 is a small effect but its significance is two-fold. It is a sequence-derived, zero-fit, fully transferable prior, no parameters are tuned to ASD, and the same alphabet applies to any protein, so the fact that a universal contact pattern carries any reproducible, correctly signed allosteric signal is the point. And, as the next section shows, the directional channels that are anti-predictive for static site annotation are in fact predictive of something else; conformational change.

### 3.4 Case studies across mechanisms

To see the sink result on individual proteins we scored four mechanistically diverse canonical systems, to span the performance range. Carbamoyl phosphate synthetase (PDB id: 1A9X)^27^, a classic allosteric enzyme, scores best (per-chain sink-AUC 0.86), with ASD sites falling directly under the tallest sink peaks. The P2X7 ion channel bound to an allosteric antagonist (PDB id: 5U1U^28^, AUC 0.82) places its antagonist-site cluster (~residues 290–305) at the highest sink peak. The lac repressor (PDB id: 1EFA^29^, AUC 0.63), a DNA-binding allosteric switch, shows moderate agreement. Phosphofructokinase (PDB id: 1PFK^30^, AUC 0.22), itself a textbook allosteric enzyme, is included as an honest failure case, where the per-chain sink track does not track the annotated sites. Per-residue sink tracks with ASD sites overlaid are shown in Figure 4; the same four proteins colored by sink score on their three-dimensional structures are shown in Figure 5, and sink-scored structures (B-factor = sink *z*) render directly in molecular viewers.

**Figure 4.**
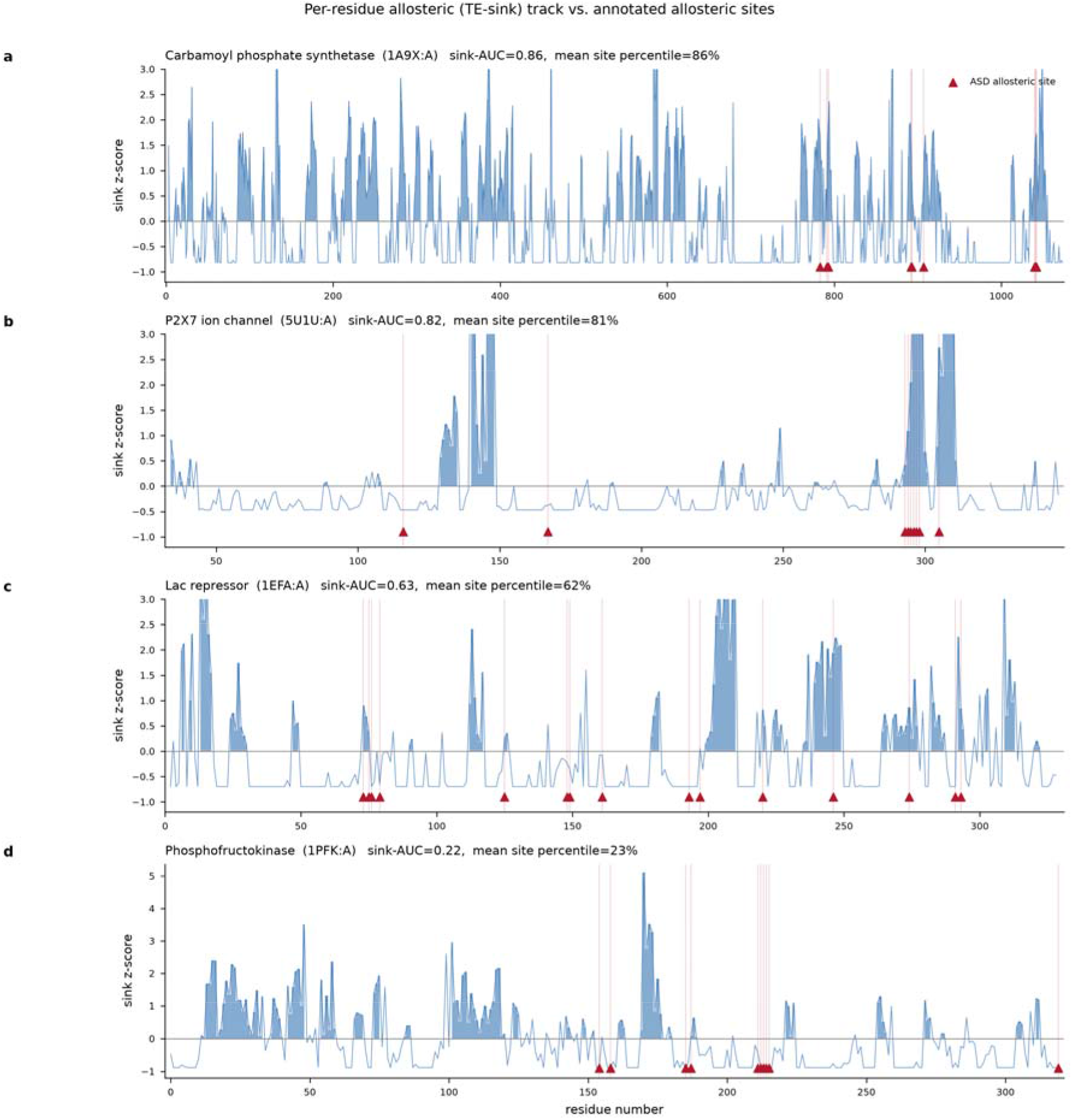
Per-residue sink track (blue) versus annotated ASD allosteric sites (red triangles) for four canonical systems spanning the performance range, including phosphofructokinase (d) as a failure case.

**Figure 5.**
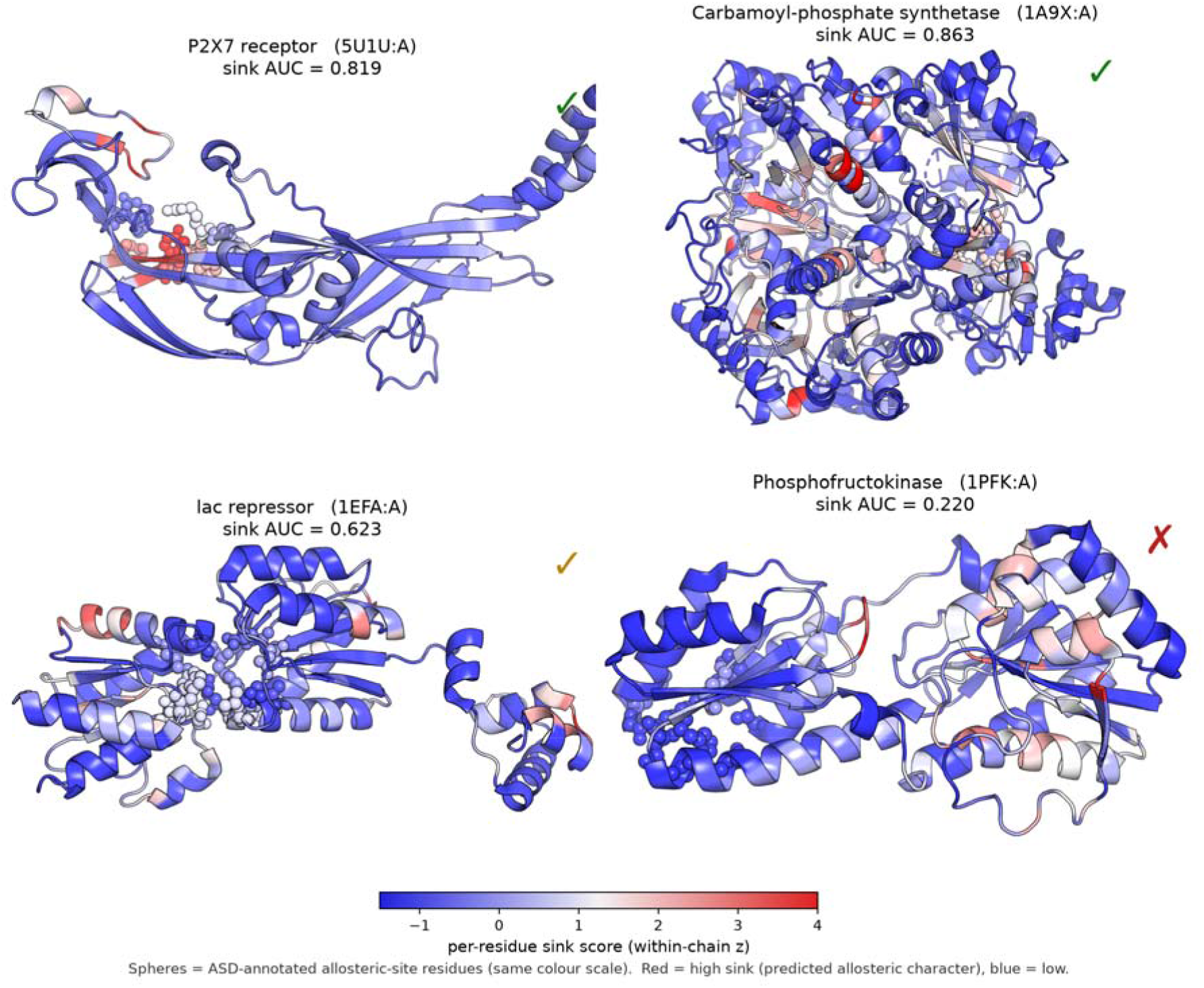
ASD allosteric sites mapped onto structure, colored by pooled alphabet sink score. Cartoon of chain A for each case-study protein, coloured by per-residue sink score (within-chain z; red = high sink, blue = low); spheres mark ASD-annotated allosteric-site residues on the same colour scale. The visual tracks the quantitative result: concordant in P2X7 (5U1U, AUC 0.82) and carbamoyl-phosphate synthetase (1A9X, 0.86), weakly positive in lac repressor (1EFA, 0.63), and the honest failure case in phosphofructokinase (1PFK, 0.22).

### 3.5 Source and sink residues play opposite roles in conformational switching

The ASD test scores a single static structure against a single static site list, and on that test the source channel is anti-predictive. But allostery is dynamic, and a natural second question is whether the directional channels predict how a protein’s information flow rewires between conformational states. We assembled nine canonical allosteric proteins that each crystallize in two well-characterized states (apo/holo and open/closed), spanning classic large-motion enzymes and signalling switches: adenylate kinase (4AKE^31^→1AKE^26^), maltose-binding protein (1OMP^32^→1ANF^33^), calmodulin ±Ca^2+^ (1CFD^34^→1CLL^35^), glucokinase (1V4T→1V4S)^36^, dihydrofolate reductase (5DFR^37^→1RX2^38^), ribose-binding protein (1URP^39^→2DRI^40^), glutamine-binding protein (1GGG^41^→1WDN^42^), citrate synthase (1CTS→2CTS)^43^, and H-Ras GDP/GTP (6Q21^44^→5P21^45^).

For each protein we computed per-residue net TE in both states with the GNM-TE method, z-normalized within each state (absolute TE scale differs between conformations), and defined the per-residue conformational rewiring as |Δ net TE| between states. We then asked which pooled alphabet channel, evaluated on the apo state alone, predicts that rewiring.

The switch channel, the pooled cross-survey TE variance, is a weak predictor (pooled Spearman ρ = +0.046). This is mechanistically sensible: switch-Std measures how variable a motif’s role is across thousands of unrelated proteins, which is not the same quantity as this protein’s state change, and it explains why the switch channel was neither a site detector (Section 3.3) nor a strong rewiring predictor. The directional channels, however, split cleanly and in opposite directions (Figure 6a, d):

**Figure 6.**
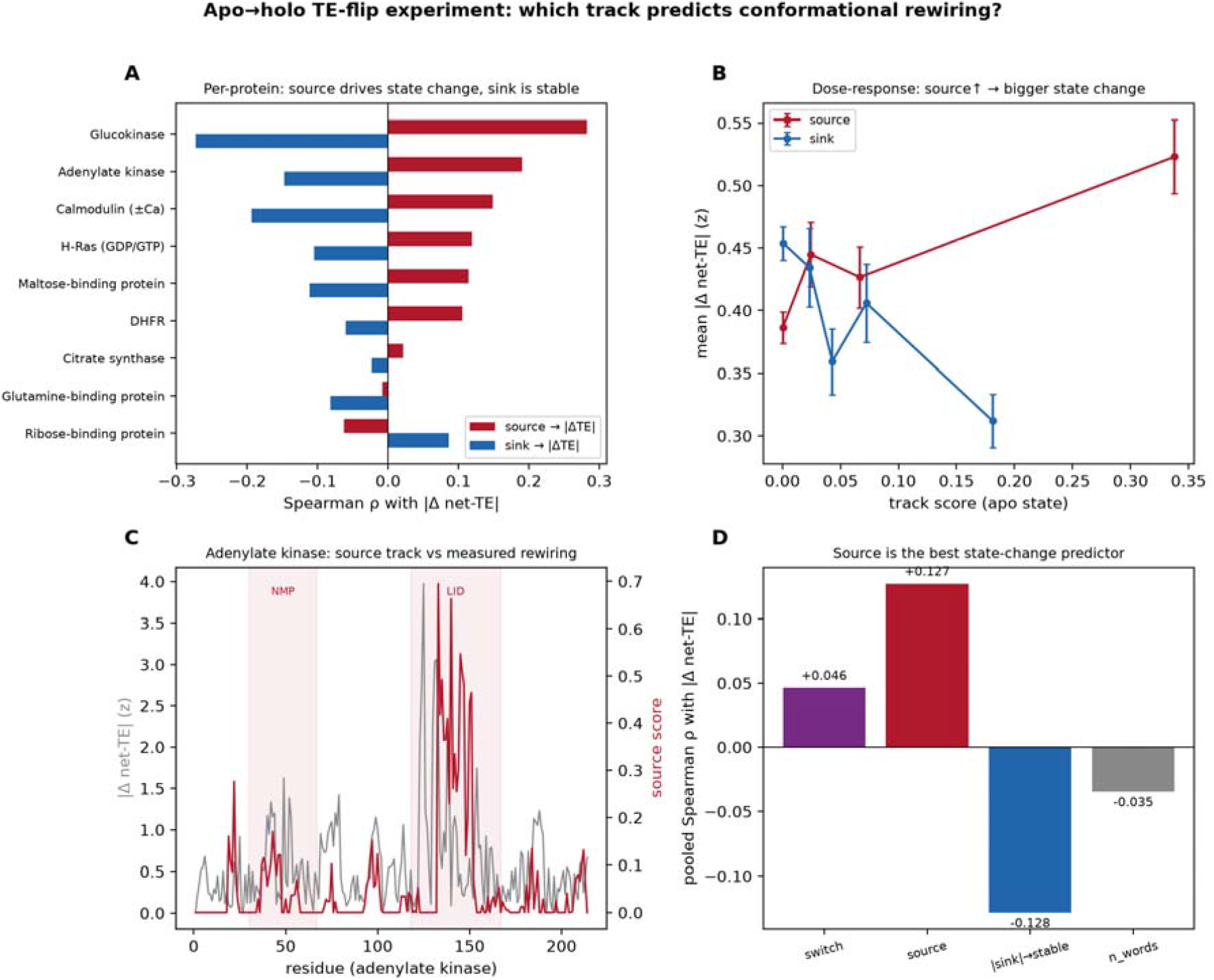
Apo → holo TE-flip experiment. Per-protein source/sink correlation with |Δ net TE| (a), pooled dose–response (b), adenylate-kinase per-residue overlay with LID/NMP marked (c), and channel comparison (d).

#### Source predicts rewiring

Source score correlates positively with |Δ | (pooled ρ = +0.127, p ≈ 4 × 10^−10^; meta-analytic ρ = +0.106, 95% CI [+0.066, +0.146]), positive in 8 of 9 proteins (Figure 6b).

#### Sink predicts stability

Sink score correlates negatively (pooled ρ = −0.128, *p* ≈ 3 × 10^-10^; meta-analytic ρ = −0.105, 95% CI [−0.145, −0.065]), negative in 8 of 9 proteins.

The strongest effects are in the textbook large-amplitude allosteric enzymes, glucokinase (ρ_source = +0.28), adenylate kinase (+0.19), calmodulin (+0.15), and in adenylate kinase the source track peaks over exactly the LID and NMP lids that clamp shut on substrate (Figure 6c). Extending this single-protein overlay to all nine two-state pairs (Figure 7) shows that the apo source track anticipates the measured conformational rewiring in 7 of 9 proteins, with the strongest concordance in glucokinase, adenylate kinase, and calmodulin.

**Figure 7.**
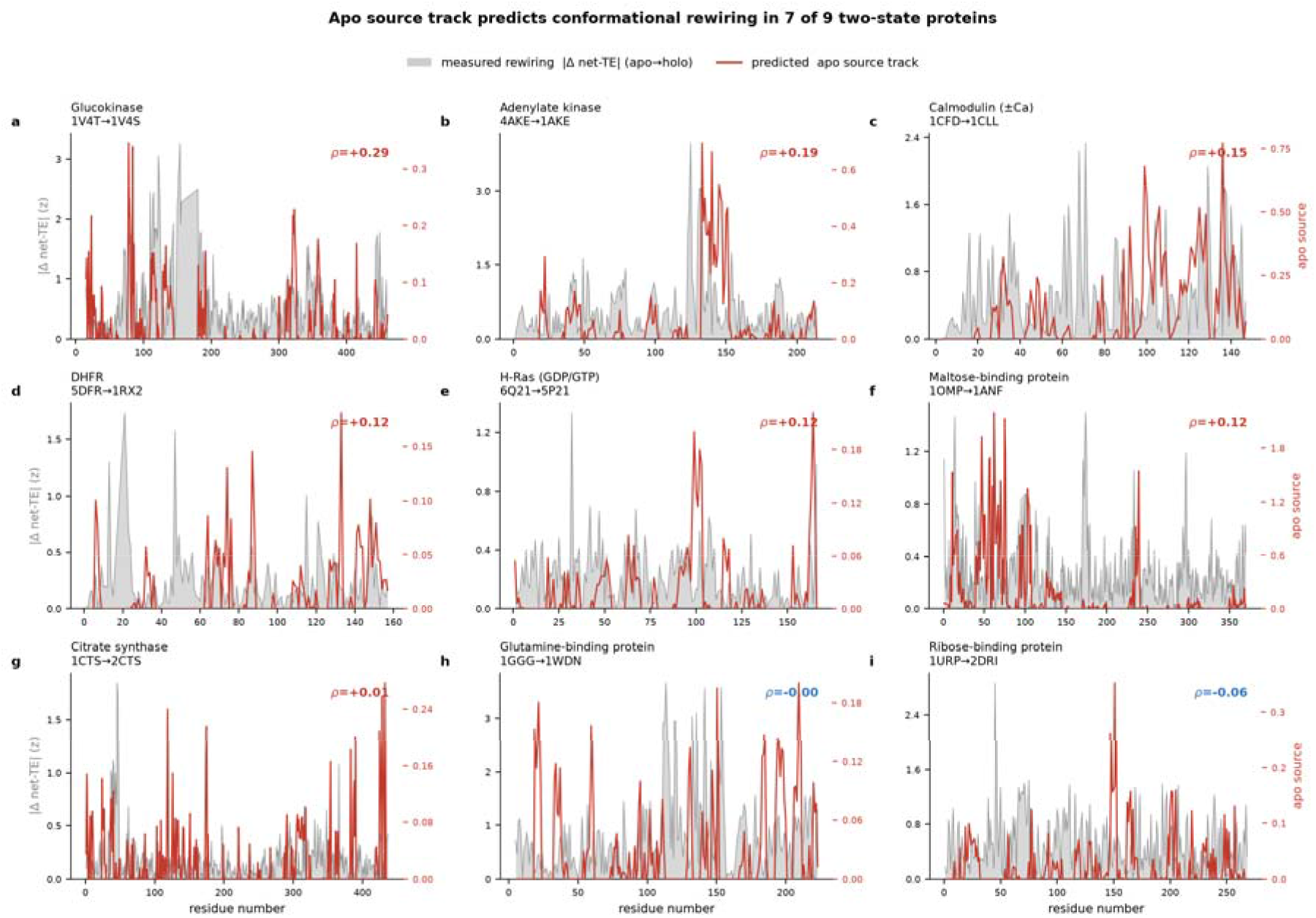
The zero-fit source track anticipates where proteins conformationally rewire. For each of nine apo → holo two-state pairs, the per-residue apo source track (red, right axis) is overlaid on the measured rewiring |Δ net TE| between the two states (grey fill, left axis). Both are z-scored within each chain; Spearman ρ(source, |Δ net-TE|) is inset per panel (red = positive, blue = negative). Panels are ordered by descending ρ. The source track is positively correlated with the actual rewiring in seven of nine proteins, strongest in the large-amplitude allosteric enzymes (glucokinase, adenylate kinase).

This resolves the apparent tension in the ASD result. Sinks mark where allosteric signal is received, binding-adjacent, conformationally *stable* anchor points that show up as ASD sites. Sources mark where allostery drives motion, the mobile machinery that rewires between states.

The directional TE decomposition thus separates the *antenna* from the *engine* and the channel that is anti-predictive for one question is the informative one for the other.

### 3.6 A switch pattern for allosteric design

Having established that the switch channel is a cross-survey context-variability measure rather than a per-protein predictor, we characterize the 21,462 high-variance switch words as a candidate design pattern, motifs whose dynamic role is unusually context-dependent, and therefore whose transplantation might reprogram coupling.

The switch set has a consistent signature:

**Source-biased**. 87% of switch words have positive mean TE (emitters), versus 50% in the background, switch motifs preferentially drive rather than receive, consistent with the source/rewiring result above.

**Compact**. Switch words have shorter linkers than background (median length 4, with a thinner long-range tail), coupling through local packing rather than long loops.

**Hydrophobic-enriched**. Clique/contact positions are enriched in bulky aliphatic hydrophobics (Leu, Ile, Met, Phe, Val; log_2_ fold-change up to +0.4) and depleted in Pro, Gly and Asp, coupling through packable hydrophobic contacts, not flexible turns.

We rank the vocabulary by a *design score (per - word TE standard deviation)* × *(log*_*10*_ *coun + 1)*, balancing context-sensitivity against statistical reliability. The top motifs include ClAS, CLaS, CLAS (sink-leaning) and RfLR, VkNE, IPpS (source-leaning). These are released as a ranked dictionary (*switch_vocabulary.csv*) of candidate transplantable coupling contacts (Figure 8).

**Figure 8.**
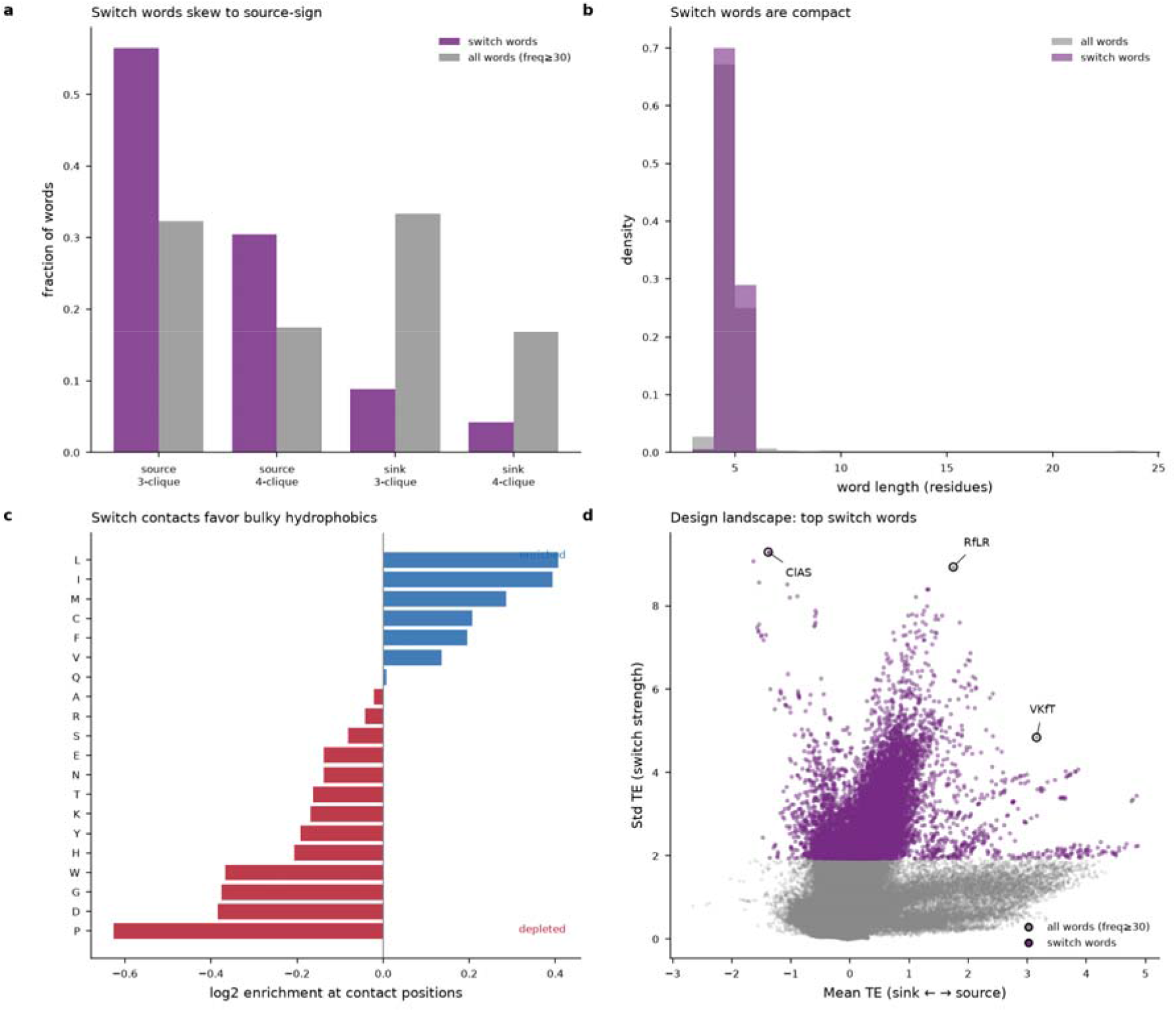
The switch design vocabulary: source-sign skew (a), compact word lengths (b), amino-acid enrichment at contact positions (c), and the ranked design landscape (d).

Although the switch-vocabulary residues are not enriched at ASD ligand-binding allosteric sites (sink is the site discriminator, Section 3.3), the per-residue switch channel is significantly elevated at documented conformational switches and hinges (Figure 9). Across four canonical allosteric proteins scored with the same zero-fit method, the switch channel is raised inside classically defined switch elements, the adenylate kinase LID domain (residues 118–167; within-region mean switch z = +0.80, one-sided Mann–Whitney p = 6×10^-10^) whose ligand-induced closure is a canonical conformational transition^46^, the calmodulin central hinge (74–82; z = +0.79, p = 0.003) which controls interdomain flexibility and target-induced conformational rearrangement^47^, and the Ras Switch I loop (30–38; z = +0.85, p = 0.006) a nucleotide-dependent signalling switch involved in effector recognition^48^. Other regions (Ras Switch II, the adenylate kinase NMP-binding hinge and both of the DHFR loops) show no enrichment. The DHFR comparison is biologically motivated because its Met20, FG, and GH loops are known to undergo functionally important fluctuations during catalysis.^49, 50^ The switch channel reports conformational switch and hinge machinery rather than ligand pockets, consistent with its near-chance performance against ASD sites and with its intended use as a design vocabulary.

**Figure 9.**
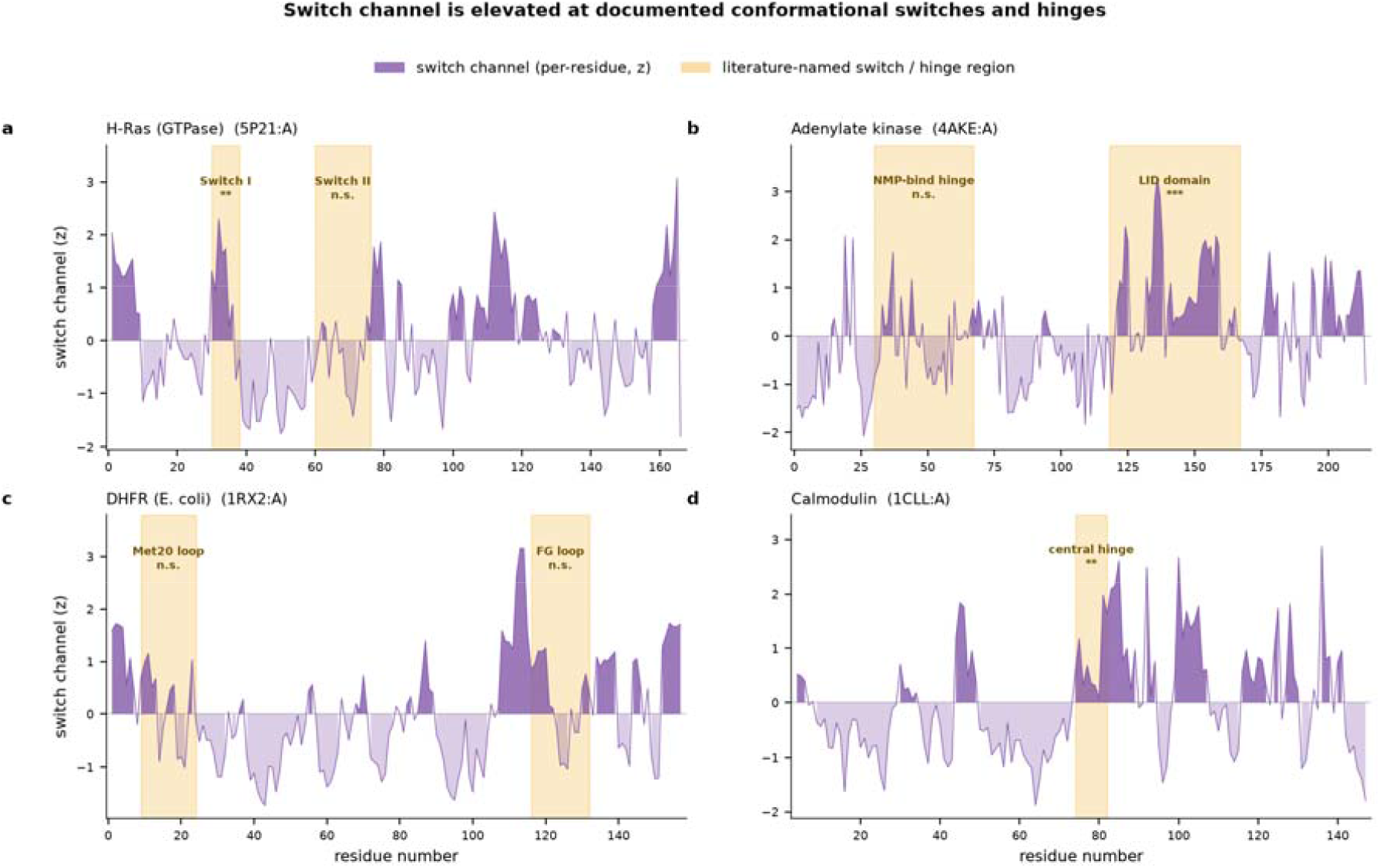
The switch channel is elevated at documented conformational switches and hinges. Per-residue switch channel (z-scored within chain, purple) for four canonical allosteric proteins, with literature-defined switch/hinge regions shaded (gold) and one-sided Mann–Whitney significance marked (*** p<0.001, ** p<0.01, * p<0.05, n.s. not significant).

## 4. Discussion

The central validated result, that ASD allosteric-site residues are transfer-entropy sinks, has a clean physical reading. Annotated allosteric sites are, operationally, *modulator-binding pockets*: the places where a regulatory ligand enters and where the dynamic consequence of binding is integrated. Such positions are information *receivers*, and they are conformationally stable enough to form a defined pocket, precisely the sink signature we recover, and precisely why the source channel is anti-predictive for this ground truth. The two-state experiment closes the loop: the same source channel that fails to mark static pockets is the one that predicts where the protein moves.

The ASD sink-AUC of ~0.54 is not a competitive site predictor and we do not present it as one. It is a *universal, zero-fit prior* built from the physics of information flow, and its value is conceptual (a transferable contact vocabulary carries real, correctly signed allosteric signal) and mechanistic (its channels decompose into receiving versus driving). Several concrete routes could raise the absolute performance: multi-chain / interface scoring (the current per-chain scan misses the many ASD sites that sit at oligomeric interfaces), pocket-neighbourhood aggregation rather than single-residue scoring, and structure-context stratification (the best-scoring case, carbamoyl phosphate synthetase at ~1,056 residues, hints that large multidomain enzymes carry more signal).

The switch dictionary is a design-stage output: compact, hydrophobic, source-biased motifs whose context-variable TE role makes them candidate coupling elements. The obvious experimental test is transplantation, grafting a top source-switch motif such as RfLR into a permissive scaffold and assaying for induced allosteric coupling. We present the vocabulary explicitly as a set of testable hypotheses, not validated designs.

GNM treats fluctuations as isotropic and captures only the topology of the contact network, so directional TE here inherits that coarse graining; anisotropic network models would refine the directional channels. ASD annotation is incomplete and conflates catalytic with allosteric sites in some entries, which bounds the achievable validation AUC. Scoring is currently per-chain rather than per-complex. The two-state experiment, while coherent across nine proteins, rests on modest per-protein correlations whose aggregate significance, not any single protein, is the claim.

## 5. Data and code availability

The scoring method (*allo_alphabet.py*), the GNM-TE core (*allo_worker.py*), the pooled pattern database (*pooled_pattern_database.parquet*), the ASD per-residue score table (*asd_residue_scores.csv.gz*), the two-state metrics (*te_flip_metrics.csv*), the switch vocabulary(*switch_vocabulary.csv)* are available in Zenodo, https://zenodo.org/records/21668707.

